## Supplementary material for "Beneficial cumulative effects of old parental age on offspring fitness": All supplementary materials

**Supplementary Methods**

**Cumulative parental effects over six generations assay**

In the multigenerational assay, lines from the parental propagation regimes (*young*, *old* and *switched*) went extinct when no eggs where produced or the eggs produced did not hatch. If no eggs where laid on day 1 in the *young* propagation regime, we attempted to collect eggs again on day two to prevent the loss of lines due to slow sexual maturation. Over the six generations, in the *young* propagation regime, we only took offspring from two-day old adults instead of one day olds on 10 occasions.

**Supplementary Figures**

**Figure S1**: Total reproduction for three parental propagation regimes across generations. Total reproduction (± SE) of offspring from *young (yellow)*, *old (purple)* and *switched (green)* parental age propagation regime after one, three and six generations.

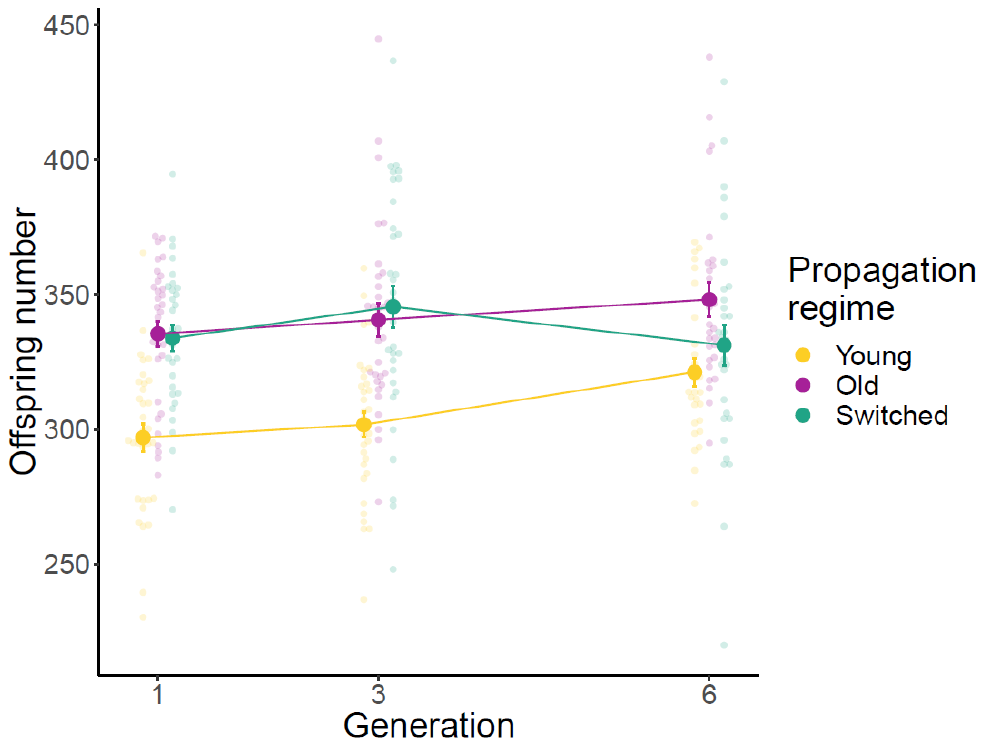

**Figure S2**: Total reproduction in generation seven. Total reproduction (± SE) of offspring from one- and three-day old parents (proximate parental age on the X axis) generated from the *young* (yellow) and *old* (purple) parental propagation regime at generation seven.

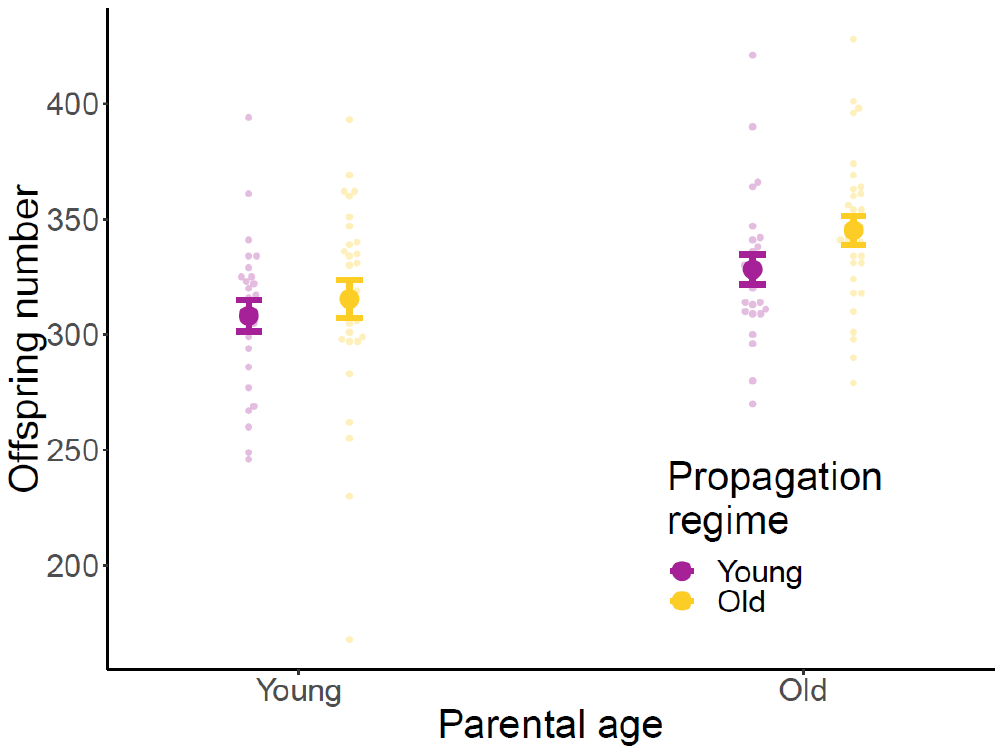

**Supplementary Tables**

| **Table S1:** Model summary for lifespan | |  |  |  |  |  |
| --- | --- | --- | --- | --- | --- | --- |
|  | Lifespan (mixed effects cox proportional hazards: N= 196) | | | | |  |
| **Fixed/*Random* effects** | **Coef** | **SE (Coef)** | **z** | **p** | **Var** | **SD** |
| Propagation regime (Old) | 0.012 | 0.189 | 0.06 | 0.95 | - | - |
| Propagation regime (Switched) | -0.056 | 0.186 | -0.3 | 0.76 | - | - |
| Generation | -0.316 | 0.096 | -3.29 | **0.001** | - | - |
| *Ancestral line* | - | - | - | - | 0 | 0.009 |
| Coefficients, standard errors, test statistics and variance components are taken from a mixed effects cox proportional hazards model on survival. Effects associated with a p-value smaller than 0.05 are highlighted in bold. | | | | | | |

| **Table S2:** Model summaries for total reproduction and lambda in generations one, three and six | | | | | | |  |  |  |  |  |  |  |  |  |
| --- | --- | --- | --- | --- | --- | --- | --- | --- | --- | --- | --- | --- | --- | --- | --- |
|  | Total Reproduction F1, 3 and 6 (LMM: N= 275) | | | |  |  |  |  | Individual Fitness (Lambda) F1, 3 and 6 (log LMM: N= 275) | | | | |  |  |
| **Fixed/*Random* effects** | **Coef** | **SE (Coef)** | **df** | **t** | **p** | **Var** | **SD** |  | **Coef** | **SE (Coef)** | **df** | **t** | **p** | **Var** | **SD** |
| Intercept (Young regime) | 290.102 | 6.444 | 269 | 45.019 | **< 0.001** | - | - |  | 1.593 | 0.022 | 260 | 71.792 | **<0.001** | - | - |
| Propagation regime (Old) | 42.868 | 9.176 | 269 | 4.672 | **< 0.001** | - | - |  | 0.092 | 0.031 | 241 | 2.918 | **<0.005** | - | - |
| Propagation regime (Switched) | 49.853 | 9.194 | 269 | 5.422 | **< 0.001** | - | - |  | 0.177 | 0.032 | 240 | 5.602 | **<0.001** | - | - |
| Generation | 4.929 | 1.741 | 269 | 2.877 | **0.004** | - | - |  | 0.018 | 0.011 | 243 | 1.662 | 0.098 | - | - |
| Propagation regime Old:Generation | -2.397 | 2.419 | 269 | -0.991 | 0.322 | - | - |  | 0.050 | 0.015 | 243 | 3.38 | **<0.001** | - | - |
| Propagation regime Switched:Generation | -5.816 | 2.393 | 269 | -2.431 | **0.016** | - | - |  | -0.020 | 0.015 | 241 | -1.381 | 0.169 | - | - |
| *Ancestral line (33 levels)* | - | - | - | - | - | 0 | 0 |  | - | - | - | - | - | 0 | 0.007 |
| Coefficients, standard errors, test statistics and variance components are taken from LMM's on the number of offspring produced (total reproduction) and individual fitness (lambda). Effects associated with a p-value smaller than 0.05 are highlighted in bold. | | | | | | | | | | | | | | | |

| **Table S3:** Pairwise comparisons of total reproduction and lambda in generations one, three and six | | | | | | | | |  |  |  |  |  |
| --- | --- | --- | --- | --- | --- | --- | --- | --- | --- | --- | --- | --- | --- |
|  | Pairwise differences total reproduction F1, 3 and 6 | | | | |  |  |  | Pairwise differences individual fitness (Lambda) F1, 3 and 6 | | | | |
| **Generation 1** | **Coef** | **SE (Coef)** | **df** | **t.ratio** | **p** |  |  |  | **Coef** | **SE (Coef)** | **df** | **t.ratio** | **p** |
| Young vs Old | -38.48 | 8.28 | 237 | -4.7 | **< 0.001** |  |  |  | -0.133 | 0.02 | 237 | -6.543 | **< 0.001** |
| Young vs Switched | -36.85 | 8.35 | 236 | -4.4 | **< 0.001** |  |  |  | -0.154 | 0.02 | 236 | -7.495 | **< 0.001** |
| Old vs Switched | 1.63 | 8.41 | 239 | 0.2 | 0.9795 |  |  |  | -0.021 | 0.02 | 239 | -0.998 | 0.5789 |
| **Generation 3** | **Coef** | **SE (Coef)** | **df** | **t.ratio** | **p** |  |  |  | **Coef** | **SE (Coef)** | **df** | **t.ratio** | **p** |
| Young vs Old | -38.89 | 8.21 | 236 | -4.7 | **< 0.001** |  |  |  | -0.21 | 0.02 | 236 | -10.39 | **< 0.001** |
| Young vs Switched | -43.73 | 8.15 | 235 | -5.4 | **< 0.001** |  |  |  | -0.14 | 0.02 | 235 | -7.003 | **< 0.001** |
| Old vs Switched | -4.84 | 8.28 | 237 | -0.6 | 0.828 |  |  |  | 0.07 | 0.02 | 237 | 3.414 | **<0.005** |
| **Generation 6** | **Coef** | **SE (Coef)** | **df** | **t.ratio** | **p** |  |  |  | **Coef** | **SE (Coef)** | **df** | **t.ratio** | **p** |
| Young vs Old | -26.92 | 8.87 | 243 | -3 | **0.0075** |  |  |  | -0.232 | 0.02 | 243 | -10.64 | **< 0.001** |
| Young vs Switched | -10.04 | 8.65 | 240 | -1.2 | 0.4778 |  |  |  | -0.112 | 0.02 | 240 | -5.238 | **< 0.001** |
| Old vs Switched | 16.88 | 8.57 | 241 | 2 | 0.1219 |  |  |  | 0.121 | 0.02 | 241 | 5.729 | **< 0.001** |
| Coefficients, standard errors, test statistics and variance components are taken from pairwise comparisons on the number of offspring produced (total reproduction) and individual fitness (lambda). Effects associated with a p-value smaller than 0.05 are highlighted in bold. | | | | | | | | | | | | | |

| **Table S4:** Model summaries for total reproduction and individual fitness (lambda) in generation seven | | | | | | |  |  |
| --- | --- | --- | --- | --- | --- | --- | --- | --- |
| **Trait** | **Fixed/*Random* effects** | **Coef** | **SE (Coef)** | **df** | **t** | **p** | **Var** | **SD** |
| Total reproduction F7 (LMM: N= 110) |  |  |  |  |  |  |  |  |
|  | Intercept (young regime:young parental age) | 305.828 | 6.403 | 98.981 | 47.766 | **<0.001** | - | - |
|  | Propagation regime (Old) | 11.58 | 6.909 | 92.791 | 1.676 | 0.097 | - | - |
|  | Parental age (Old) | 25.475 | 6.799 | 80.268 | 3.747 | **<0.001** | - | - |
|  | *Ancestral line (33 levels)* | - | - | - | - | - | 109.8 | 10.48 |
| Individual Fitness (λind) F7 (LMM: N= 110) |  |  |  |  |  |  |  |  |
|  | Intercept (young regime:young parental age) | 3.655 | 0.034 | 107 | 108.353 | **<0.001** | - | - |
|  | Propagation regime (Old) | 0.022 | 0.038 | 107 | 0.584 | 0.56 | - | - |
|  | Parental age (Old) | 0.202 | 0.038 | 107 | 5.351 | **<0.001** | - | - |
|  | *Ancestral line (33 levels)* | - | - | - | - | - | 0 | 0 |
| Coefficients, standard errors, test statistics and variance components are taken from LMM's on the number of offspring produced (total reproduction) and individual fitness (lambda). Effects associated with a p-value smaller than 0.05 are highlighted in bold. | | | | | | | | |

| **Table S5:** Model summaries for egg size, development time and adult size | | | |  |  |  |  |  |
| --- | --- | --- | --- | --- | --- | --- | --- | --- |
| **Trait** | **Fixed/*Random* effects** | **Coef** | **SE (Coef)** | **df** | **t** | **p** | **Var** | **SD** |
| Egg size (LMM: N= 283) |  |  |  |  |  |  |  |  |
|  | Intercept | 1.096 | 0.026381 | 2.292 | 41.54 | **<0.001** | - | - |
|  | Parental age | 0.07877 | 0.009677 | 240.751 | 8.14 | **<0.001** | - | - |
|  | *Parent (70 levels)* | - | - | - | - | - | 0 | 0.034 |
|  | *Block (2 levels)* | - | - | - | - | - | 0 | 0.029 |
| Development to sexual maturity latency (LMM: N=54) |  |  |  |  |  |  |  |  |
|  | Intercept | 2.577 | 0.026 | 38.513 | 98.531 | **<0.001** | - | - |
|  | Parental age | -0.045 | 0.016 | 26.000 | -2.877 | **0.008** | - | - |
|  | *Parent (27 levels)* | - | - | - | - | - | 0.002 | 0.046 |
| Adult size at sexual maturity (LMM: N= 94) |  |  |  |  |  |  |  |  |
|  | Intercept | 0.095 | 0.002 | 57.623 | 60.642 | 60.642 | - | - |
|  | Parental age | 0.006 | 0.001 | 46.000 | 5.717 | **<0.001** | - | - |
|  | *Parent (47 levels)* | - | - | - | - | - | 0.000 | 0.001 |
| Coefficients, standard errors, test statistics and variance components are taken from LMM's on egg size, development time and adult body size. Effects associated with a p-value smaller than 0.05 are highlighted in bold. | | | | | | | | |
